# An in-depth analysis and exploreation with focus on the biofilm in *Staphylococcus aureus*

**DOI:** 10.1101/2024.05.05.592613

**Authors:** Zhiyuan Zhang, Guozhong Chen, Yuanyuan Pan, Zhu Yang, Yin Liu, Erguang Li

## Abstract

Research into the biolfilm formation in *Staphylococcus aureus* has benefited greatly from the generation of high-throughput sequencing data to drive molecular analyses. The accumulation of these data, particularly transcriptomic data, offers a unique opportunity to unearth the network and constituent genes involved in the biofilm formation of *Staphylococcus aureus* through machine learning strategies and co-expression analyses. Herein, we harnessed all available RNA sequencing data related to *Staphylococcus aureus* biofilm studies and identified influenced functional pathways and corresponding genes in the process of the transition of bacteria from planktonic to biofilm state via employing machine learning and differential expression analyses. By weighted gene co-expression analysis and our previously developed predictor, important functional modules, potential biofilm-associated proteins and subnetwork of biofilm formation pathway were found. By constructing a protein-protein interaction (PPI) network, we discovered several hitherto unreported novel protein interactions within these functional modules. To make these data more straightforward to experimental biologists, an online database named SAdb was developed (http://sadb.biownmcli.info/).

**IMPORTANCE:** In this work, we conducted a comprehensive and in-depth exploration of RNA sequencing data in biofilm research through differential expression analysis, machine learning, WGCNA, and biofilm-associated protein predictive analysis, which has also illuminated novel analytical perspective for other research into bacterial phenotypes. And, to provide researchers with unimpeded access to these data, we developed a database name SAdb for the storage and analysis of *Staphylococcus aureus* omics data. We believe that this study will captivate the interest of researchers in the field of bacteriology, particularly those studying biofilms, which play a crucial role in bacterial growth, pathogenicity, and drug resistance.

## 1. Introduction

*Staphylococcus aureus*, a ubiquitous pathogen, is implicated in a myriad of afflictions ranging from pneumonia and sepsis to infections of wound and the urinary tract (1). The capacity of pathogen to form bacterial biofilm during the infectious cycle poses a formidable challenge to medical intervention (2–3). it is documented that biofilm-ensconced bacteria exhibit approximately a thousandfold increased resistance to antimicrobial agents compared to their planktonic counterparts (4). Additionally, a significant proportion of all microbial infections—65%, and chronic infections—80%, respectively, are attributed to biofilm genesis (5). Thus, elucidating the biofilm development mechanisms of *Staphylococcus aureus* is imperative for enhancing treatment efficacy and providing a theoretical framework for its prevention and management.

The bacterial biofilm is an intricate membrane-like structure constituted by extracellular DNA (eDNA), polysaccharides, various proteins, and other bacterial secretions (6). Among these, biofilm-associated proteins constitute a vital category that includes both structural components of the biofilm and regulatory elements governing its formation. The exploration and identification of these proteins are crucial for understanding the molecular intricacies of bacterial biofilm formation. Typically, these proteins are identified through experiments where, for instance, proteins such as CcpA, CodY, and GltS have been determined to influence the formation of biofilms in *Staphylococcus aureus* (7–8). Moreover, when compared to organisms like *E. coli,* whose biofilm pathway documented in the KEGG database encompasses 110 proteins, and *Vibrio cholerae that* the corresponding biofilm network includes 75 proteins, the biofilm networks of *Staphylococcus aureus* have yet to be annotated in prevalent databases, highlighting a substantial gap in our knowledge and underscoring the potential for groundbreaking research in this domain. Alongside advances in omics research, the *Staphylococcus aureus* have benefited greatly from next-generation sequencing (NGS) technologies (9). The power of meta-analysis has been amplified with the integration of high-quality bacterial NGS data, allowing for more robust conclusions from varied experimental results. Particularly, expression data lend themselves well to such integrative analysis, facilitating the categorization of genes into functional modules through co-expression metrics. In microbial research, methodologies such as the weighted gene co-expression network analysis (WGCNA) have proved invaluable in delineating critical genetic interactions under the "guilt by association" principle (10). Moreover, machine learning techniques continue to reveal new biological insights by analyzing vast and varied datasets, particularly microarray and sequencing data (11).

In our current endeavor, we had amalgamated RNA sequencing data from *Staphylococcus aureus* under biofilm phenotype and analyzed the data using differential expression analysis, machine learning, weighted correlation network analysis (WGCNA), and our developed predictor (http://124.222.145.44/#!/score). We hope to identify important proteins and their interactions under the biofilm phenotype through a comprehensive analysis strategy. Additionally, to facilitate researchers ’ easy access to our data, we have developed an online database named SAdb, which also collects, organizes *Staphylococcus aureus* gene expression data, gene annotation information, and proteomic data, available publicly at http://sadb.biownmcli.info/.

## 2. Results

### 2.1 Analysis procedures

For initial raw data collection, we downloaded and curated four ***Staphylococcus aureus*** transcriptome (RNA-seq) datasets in biofilm research, in total 171 samples, from the European Nucleotide Archive (EBI ENA, https://www.ebi.ac.uk/ena/browser/home) (12) and Sequence Read Archive (SRA, https://www.ncbi.nlm.nih.gov/sra/) (13). For the processing of raw sequencing reads, FastQC (http://www.bioinformatics.babraham.ac.uk/projects/fastqc/) was used to evaluate the overall quality of the raw sequencing reads, followed by the Trim_galore to remove sequencing vectors and low-quality bases (14) and performed transcript quantification using Salmon (15), which adopted TPM (Transcript Per Million) for normalization, a better unit for RNA abundance than RPKM and FPKM since it respects the invariance property and is proportional to the average relative RNA molar concentration (16). All RNA-seq samples were mapped to a recently inferred ***Staphylococcus aureus*** transcriptome derived from the strain NCTC 8325 reference genome. We integrated data from four datasets via the Sva R package Combat function and performed differential expressed analysis, in which we used a cutoff of |log2 FC| > 1.0 (FC, fold change) and p-value <0.05 to define differentially expressed genes between experiments. Subsequently, we utilized the DEGs obtained for the construction of machine learning models, and through the feature gene selection strategy, filter many DEGs to identify feature genes that make significant contributions to distinguishing the biofilm phenotype and planktonic state under two experimental conditions. We analyzed these feature genes using the biofilm-associated protein predictor previously developed. Finally, we used the WGCNA (https://github.com/ShawnWx2019/WGCNA-shinyApp) pipeline to analyze the entire gene expression table and generate a co-expression network. The workflow of analysis is shown in **Fig. 1**.

**Fig. 1.**
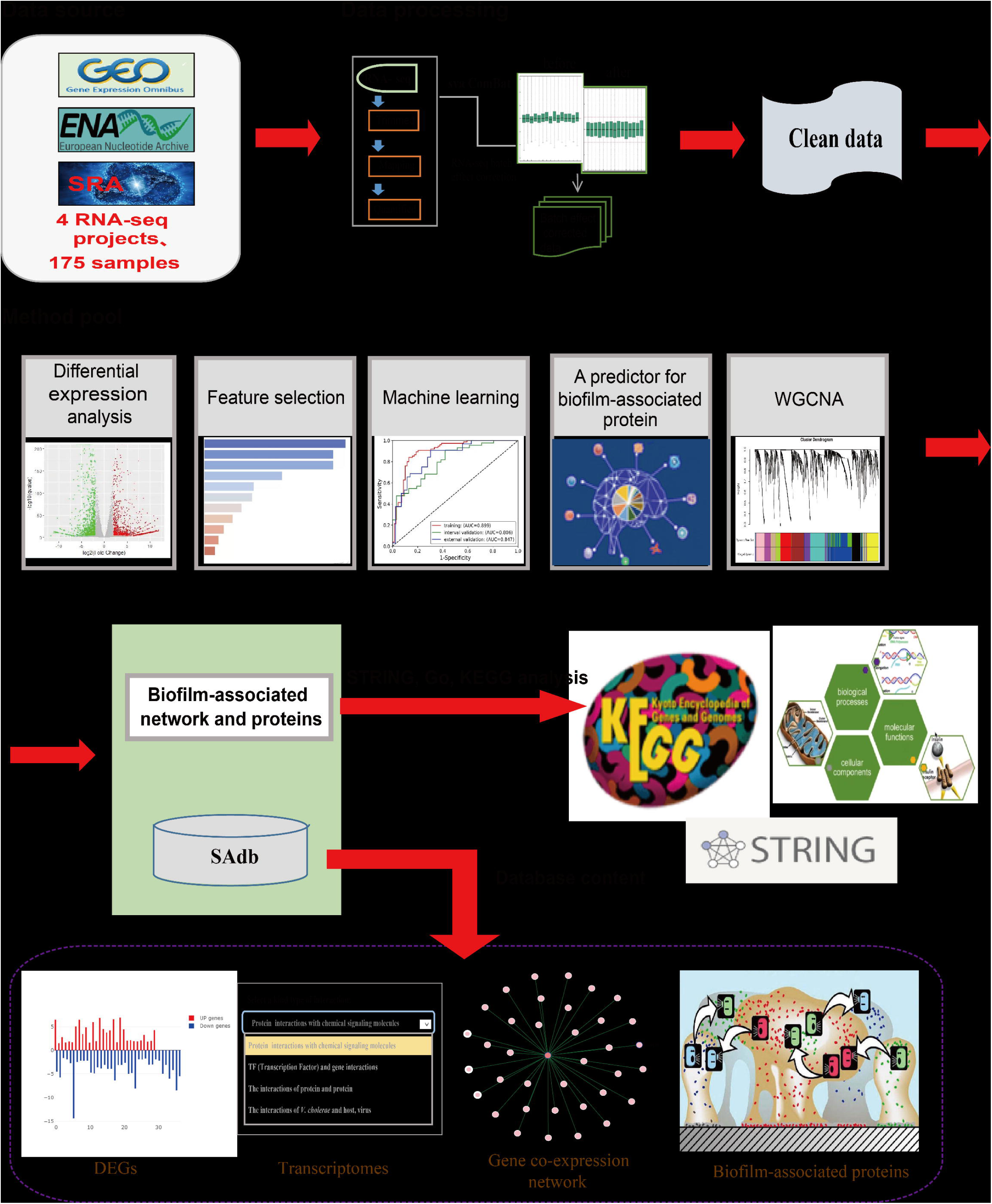
The workflow of analysis. RNA sequencing data were collected from public databases, including GEO, ENA and SRA. Then, The RNA sequencing raw data perform quality control, mapping, transcript quantification and normalization to obtain high-quality data for further analysis. Next, biofilm-associated network and proteins were obtained via differential expressed analysis, machine learning, prediction analysis and WGCNA. Finally, comprehensice database named SAdb was developed.

**Fig. 2.**
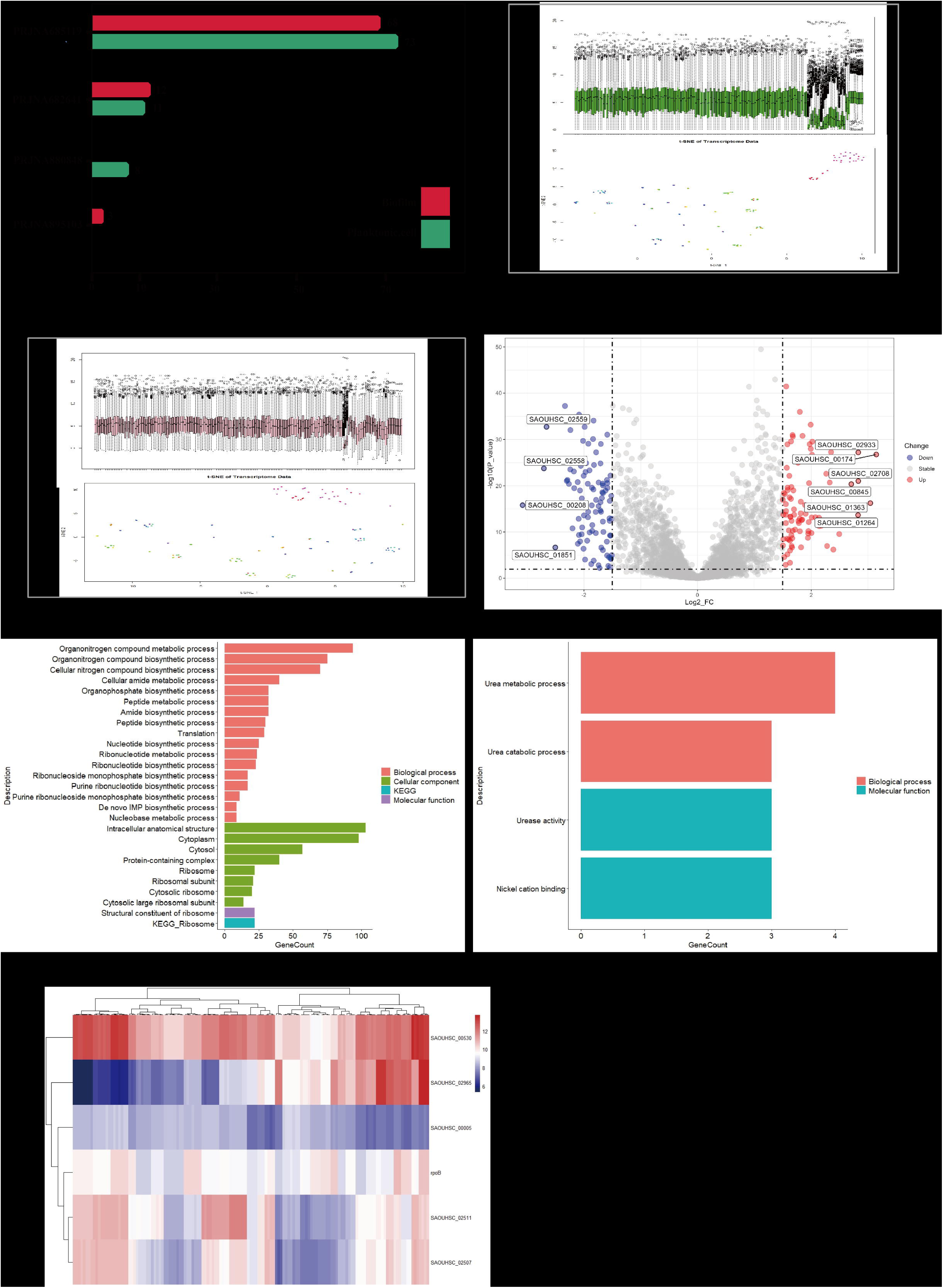
The acquisition of differentially expressed genes and the corresponding GO and KEGG analyses. (A) The source and size of RNA sequencing data, which also provides the size of experimental samples. (B) The deployment of box plots and t-SNE analyses to exhibit batch effects among different datasets. (C) The employment of the sva Combat function to exterminate batch-related discrepancies, aiming to procure a dataset of elevated quality. (D) The utilization of volcano plots to showcase differentially expressed genes that meet the specified criteria (|Log2FC| >1, p-value < 0.01). (E) GO and KEGG analyses on the DEGs to identify enriched functional pathways. (F) The selection of the top 5 upregulated and downregulated genes for subsequent GO and KEGG analyses to determine the enrichment pathways. (G) The assessment of the utility of internal reference genes within Staphylococcus aureus.

### 3.2 Data integrating and analysis

Through boxplot analysis and the R t-SNE function, we found that there was a batch effect in the expression profile between different batches of items from RNA-seq. We normalized these data using the Sva R package Combat function. The boxplot results show that the gene expression profiles of different samples are more consistent compared to before. These results indicate that we have integrated data originating from different datasets, which can be used for further analysis. Next, we first examined the transcriptional profiles of *Staphylococcus aureus* samples and obtained 409 DEGs through differential expression analysis, accounting for 14 percent of the total number of genes, which indicates that multiple pathways of *Staphylococcus aureus* are affected by environmental changes.

The subsequent functional enrichment, facilitated by the Gene Ontology (GO) (17) and Kyoto Encyclopedia of Genes and Genomes (KEGG) (18) databases, gleaned significant insights into the impacted biological functions. The GO analysis (Fig. 3B) primarily spotlighted aspects such as biosynthesis, cytoplasm, and ribosomal structure —key indicators of growth, intracellular activities, and transcriptional mechanisms in *Staphylococcus aureus*. In a similar vein, KEGG enrichment analysis (Fig. 3C) pinpointed the ribosome as the most critical pathway. Collectively, both GO and KEGG enrichment analyses elucidate the swift transcriptional and translational responses of bacterial genes and proteins to phenotypic alterations.

**Fig. 3.**
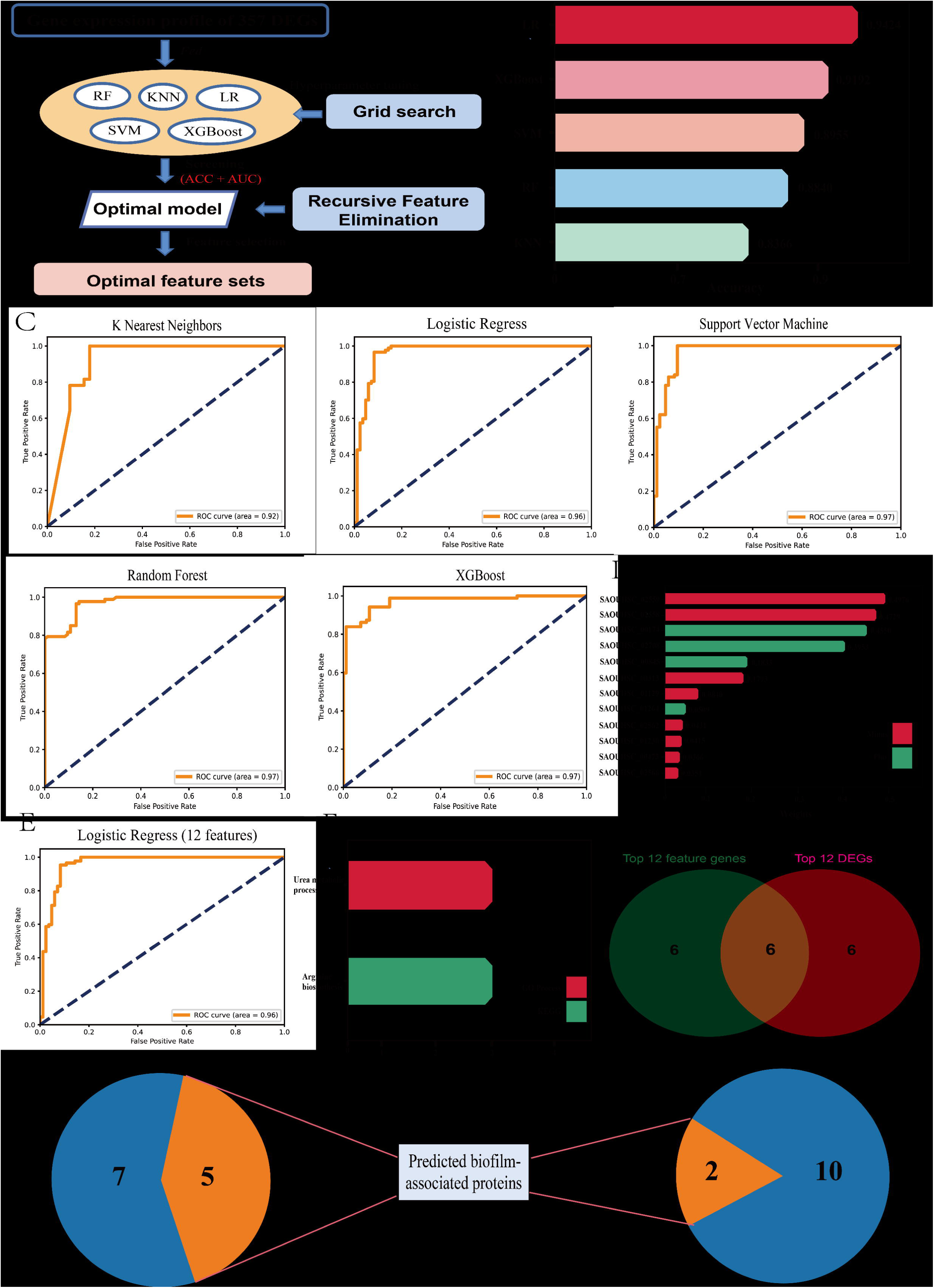
Using machine learning analysis to explore feature genes. The schematic of machine learning analysis. incorporating differentially expressed genes as data features for training machine learning models is presented, noting that genes encoding non-coding proteins within DEGs have been filtered out to refine feature genes for model training. Grid search and RFE are utilized for the determination of hyperparameters and optimal feature subsets, respectively. (B) The evaluation of performance amongst diverse machine learning algorithms through accuracy comparison. (C) The efficiency of different machine learning algorithms, assessed by comparing the area under the ROC curve. (D) The computation of contribution scores of feature genes to ascertain the optimal feature subset, culminating in a subset encompassed of 12 feature genes. (E) The AUC value of the optimal feature subset under the LR model. (F) The pathway enrichment outcomes for the subset composed of 12 genes. (G) The overlap analysis between feature genes and top DEGs. (H) The prediction for feature genes and top DEGs, utilizing biofilm-associated protein predictor.

**Fig. 4.**
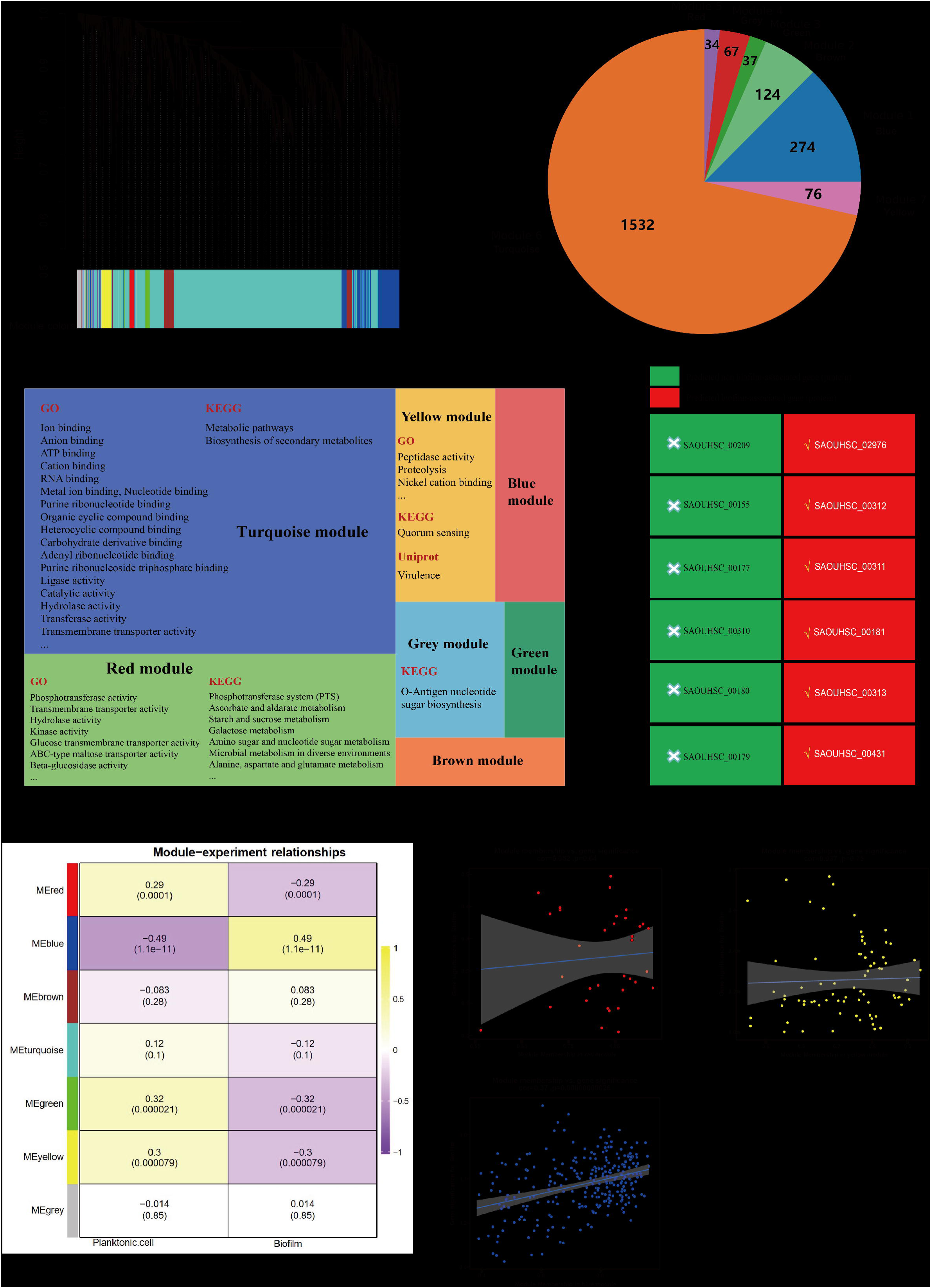
WGCNA elucidated the main functions of modules. (A) the construction of a hierarchical clustering tree based on gene co-expression analysis. (B) The size of gene modules within WGCNA and the respective gene members contained within these modules. (C) The dendrogram displaying the functional pathways enriched by gene modules. (D) The prediction of 12 feature genes using biofilm-associated protein predictor. (E) The analysis of the association between gene modules and experimental conditions, including planktonic and biofilm state. (F) The correlation analysis of genes within red, yellow, and blue modules in relation to experimental conditions.

**Fig. 5.**
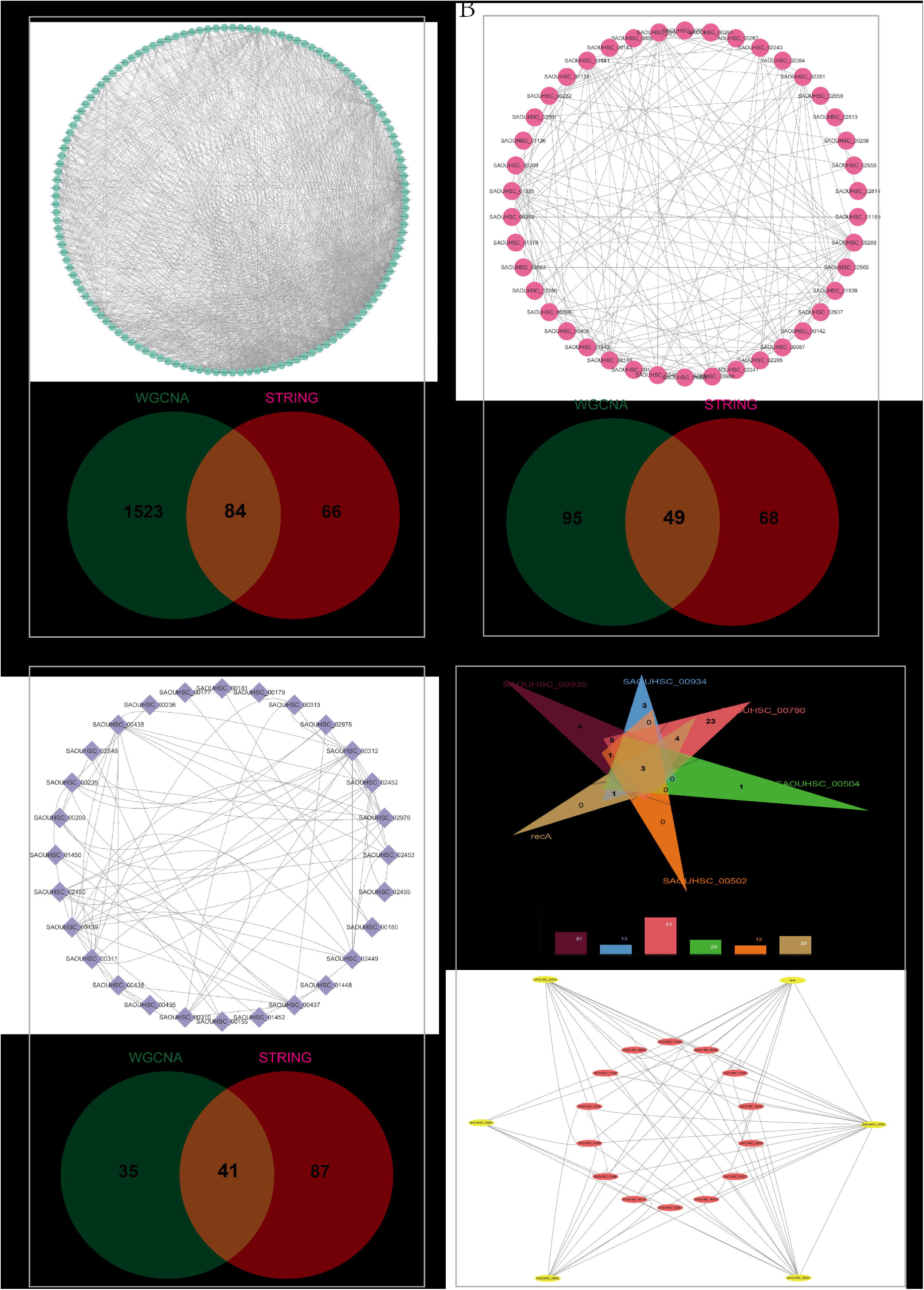
PPI interaction networks between hub genes suggest novel interactions. The interaction analysis of hub genes within the blue module and the overlapping analysis of interactions obtained through STRING database and WGCNA. (B) The interaction analysis of yellow module hub genes with an overlapping analysis of interactions derived from both STRING database and WGCNA. (C) The interaction analysis of hub genes within the red module and the overlapping analysis of interactions obtained through STRING database and WGCNA. (D) The elucidation of six experimentally validated genes encoding biofilm-associated proteins and their respective PPI networks, with an overlapping analysis to discern shared genes and potentially unveil a subnetwork within the biofilm formation pathway.

**Fig. 5.**
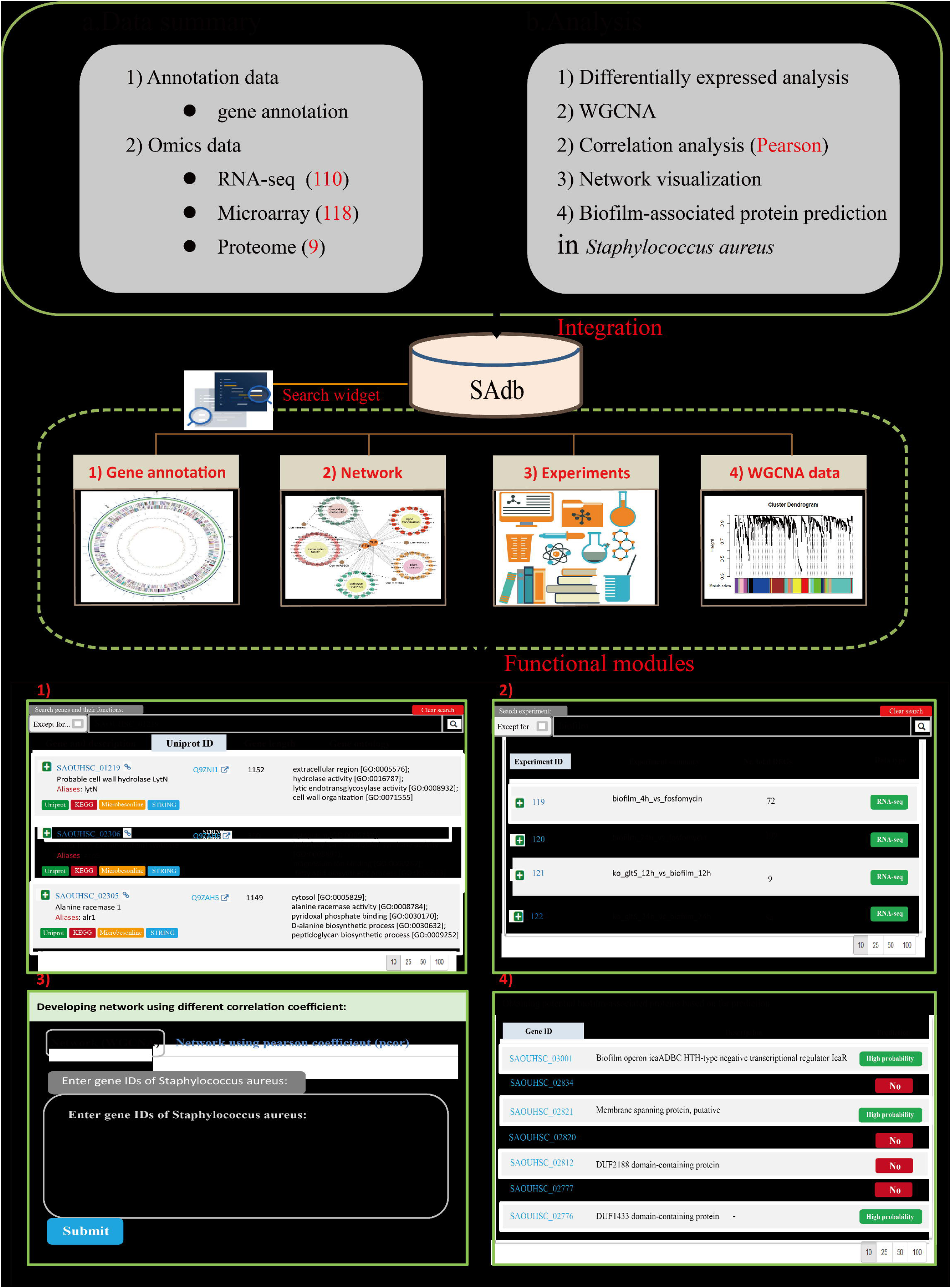
Overview of SAdb. (A) SAdb furnishing omics data along with corresponding analysis functionalities is described. (B) SAdb database includes "Genes", "Network", "Experiments", and "Results". (C) each of functional modules in SAdb is visually demonstrated.

Further exploration was directed at the principal DEGs evident during the transition between biofilm and motile states of the bacteria. The quintet of genes that were most notably downregulated at the whole transcriptome level included *SAOUHSC_02558* (*ureA*), *SAOUHSC_02559* (*ureB*), associated with nitrogen metabolism and urea degradation, alongside *SAOUHSC_02493,* a 50S ribosomal protein implicated in *Staphylococcus aureus* translation, as identified in previous research (19). *SAOUHSC_00208* and *SAOUHSC_00264* were denoted as coding for hypothetical proteins. Additionally, prominent genes such as *SAOUHSC_02561*(*ureC*), *SAOUHSC_02562* (*ureE*), which are annotated respectively as urease subunit alpha and urease accessory protein, and partake in the urea catabolic process within the KEGG pathway, with their functions in *Staphylococcus aureus* remaining unreported. Internal reference genes are very important for transcriptional-level studies, and a good internal reference gene needs to meet a certain amount of expression and stable expression. The internal reference genes selected for *Staphylococcus aureus* transcription were *gyrB* (*SAOUHSC_00005*) (20), *tuf* (*SAOUHSC_00530*) (21), *rplD* (*SAOUHSC_02511*) (22), *rpoB* (23), *rplV*(*SAOUHSC_02507*) (24), and *SAOUHSC_02965*. We found that the expression levels of *SAOUHSC_02965*, *SAOUHSC_02507*, and *rplD* were not stable under different conditions (Fig. 3D).

These genes exhibit varied expression under different experimental conditions. Choosing them as internal reference genes is risky. Relatively speaking, *SAOUHSC_00530*, *SAOUHSC_00005*, and *rpoB* are better choices as internal reference genes.

### 3.3 Using Machine learning analysis to explore feature genes

The utilization of machine learning in the domain of disease diagnostics encompasses a vast array of applications, ranging from the analysis of imagery to the intricacies of genomics, facilitating the early detection of diseases and forecasting the efficacy of treatments (25–26). Nonetheless, the exploration into bacterial phenotype via machine learning methodologies remains relatively uncharted. In this study, our endeavor is to employ machine learning algorithms coupled with strategies for feature selection to pinpoint crucial pathways that undergo modification as bacteria metamorphose from a motile state into biofilm formations, encapsulating processes such as oxidative respiration, metabolic activities, and biofilm formation. Initially, we employed expression data from 357 DEGs for the machine learning modeling datasets, as genes showing no significant changes in expression levels under two distinct experimental conditions (mobile vs. biofilm) offer no substantial contribution to distinguishing the experimental conditions, rendering it impossible to infer the corresponding experimental conditions based solely on these genes’ expression levels. Furthermore, from a machine learning model construction perspective, the inclusion of genes without significant expression variation as features augments the complexity of the model, predisposing it to overfitting. Subsequent to establishing these data inputs, we conducted a comparative analysis of model performances deploying algorithms such as Logistic Regression (LR) , XGBoost, Support Vector Machines (SVM), K-Nearest Neighbors (KNN), and Random Forest (RF), leveraging ten-fold cross-validation and optimizing hyperparameters via grid search. The findings unveiled model accuracy spanning from 83% to 94%, with Area Under the Curve (AUC) scores ranging from 0.92 to 0.97. The model devised using the LR algorithm was distinguished by achieving the pinnacle of accuracy, 94.24%, accompanied by an AUC score of 0.97, thereby selecting the LR algorithm for further feature selection and model refinement.

Recursive Feature Elimination (RFE) stands out as a feature selection procedure that iteratively reduces the number of features (27). In our study, an optimal subset of 12 genes was ascertained through the application of the recursive feature elimination approach, their contributions and significance to the model meticulously evaluated. These genes are instrumental in the bacterial transition from motility to biofilm states. The tenfold cross-validation ascertained that the LR model, when refined to include these 12 feature genes, achieved an accuracy of 93.07% with an AUC of 0.96, nearly identical to the LR model constructed with 357 DEGs. This highlights that the majority of DEGs do not play a pivotal role as features in the construction of the model. Gene Ontology (GO) and Kyoto Encyclopedia of Genes and Genomes (KEGG) analyses conducted on the 12 genes revealed an enrichment in pathways germane to urea metabolism and arginine biosynthesis, elucidating that during the transition between these two states, the bacteria exhibit heightened vigor in growth, metabolism, and the utilization of nitrogen sources. Remarkably, a third of these genes lacked functional annotations. Moreover, selecting DEGs based on log2 FC and p-value, the uppermost 12 genes also showcased enrichment in the urea metabolism pathway, reinforcing the pivotal role of urea metabolism in the bacteria’s transformation process. A subsequent overlapping analysis between the genes deemed as features and the top 12 DEGs uncovered six genes common to both sets, primarily enriched in the urea metabolism pathway, suggesting a degree of concordance in the conclusions derived through both methodologies. Intriguingly, these genes were not found to be enriched in pathways directly associated with biofilm development. Therefore, we delved into the pathway datasets of *Staphylococcus aureus* within the GO and KEGG databases, discovering an absence of annotations related to biofilm pathways, indicative of a limited understanding among researchers regarding the biofilm formation pathways by *Staphylococcus aureus.* To elucidate the linkage between these genes and the biofilm regulatory network, we analyzed the proteins encoded by these genes utilizing a predictor of bacterial biofilm-associated protein developed earlier (28). The analysis identified that among the 12 feature genes, five were prognosticated to be associated with biofilm formation, among which three are hypothetical proteins. The gene *SAOUHSC_02562* (*ureE*), an urease accessory protein, was involved in the stability of biofilms in *Staphylococcus aureus* (29). *SAOUHSC_02566* (*sarR*) has been recognized as an HTH-type transcriptional regulator, critical to the biofilm formation of *Staphylococcus epidermidis* (30). Among 12 DEGs, the hypothetical protein *SAOUHSC_00208* and a Type VII secretion system extracellular protein *SAOUHSC_00264* were predicted as potential biofilm-associated proteins.

In this study, the method of feature gene selection via machine learning, compared to the method of TOP DEGs, could enrich functional pathways that change during condition transitions while also uncovering potential biofilm-associated proteins with the predictor. However, in terms of numerical yield, the machine learning approach unravels a greater number of proteins, intimating that, in the context of handling expansive gene expression datasets, the strategy of feature gene selection via machine learning may prove more efficacious in unearthing important genes.

### 3.4 WGCNA elucidated the main functions of modules

WGCNA is a bioinformatics analysis method that reveals the interactions between gene modules and the associations between these modules and external phenotypic traits by constructing co-expression networks among genes (31). In this study, the co-expressed gene network in *Staphylococcus aureus* transcripts was revealed by the WGCNA pipeline, which distributed the filtered genes into 7 modules, with the number of genes in each module being 274, 124, 37, 67, 34, 76, 1532 respectively (Table 1). The Turquoise module, the most populous, includes 1,532 genes, half of which (50%) remain unannotated.

The Turquoise module contains 1,532 gene members, of which 766 (50%) are currently unannotated. Functional enrichment analysis of the Turquoise module mainly focused on cellular metabolism, biosynthesis, and membrane transport pathways. The Yellow module is related to bacterial virulence and quorum sensing. The Grey module’s enrichment predominantly revolves around O-antigen nucleotide sugar biosynthesis pathways. The Red module is significantly enriched in the Phosphotransferase system (PTS), Ascorbate and aldarate metabolism, and Amino sugar and nucleotide sugar metabolism. The Blue, Brown, and Green modules did not exhibit enrichment in specific functional pathways, which may be attributed to the high presence of unannotated or non-protein coding genes, with respective unannotated gene counts of 196 (71.5%), 105 (84.7%), and 28 (75.7%). In this study, we aimed to identify gene modules closely related to the biofilm phenotype, to this end, we performed an analysis of the association between experimental conditions and gene modules. The results showed that the biofilm phenotype is negatively correlated with the Red (-0.29, 0.0001), Green (-0.32, 0.000021), and Yellow (-0.3, 0.000079) modules, and positively correlated with the Blue module (0.49, 1.1e-11), indicating that when bacteria transition from mobile state to biofilm state, their virulence regulation system and energy metabolism pathways, including vitamin, sugar acid, and urea metabolism, are affected. It’s noteworthy that 54.1% of the gene members in the Green module encode tRNA and rRNA, with 21.6% remaining uncharted, leading to discontinuation in the exploration of its functionalities. Subsequently, we conducted hub gene screening analysis on the Red, Yellow, and Blue modules to enhance the understanding of the functionality expressed by these modules. The hub genes of the three modules were obtained by screening with |gene significance (GS)| > 0.2 and |module membership (MM)| > 0.8, and were analyzed for functional enrichment again. The results showed that in the Red module, 4 hub genes are enriched in Ascorbate and aldarate metabolism, 6 hub genes in the PTS system, and 4 hub genes in Amino sugar and nucleotide sugar metabolism. For unidentified gene members within this module, such as SAOUHSC_00313, the strategy of analyzing unknown genes within the same module via hub genes located in known pathways proved valuable. Moreover, research has shown that the PTS (phosphotransferase system) is related to biofilm formation (32–33). Analysis of this module’s hub genes through a predictor disclosed that 6 (50%) genes encode biofilm-associated proteins, suggesting this module may serve as a subnetwork within the biofilm regulatory pathway. In the Yellow module, four (33.4%) hub genes are enriched in the Type VII secretion system (T7SS) functional pathway, known for secreting pathogenic factors that facilitate bacterial invasion of host cells and disrupt their normal functions, thus augmenting the survival and proliferation of pathogen (34). Furthermore, T7SS aids bacteria in resource competition against other microbes (35). The secreted proteins by T7SS can directly or indirectly curtail the growth of competitors, enhancing bacterial competitiveness and survival. Regarding the hub gene analysis of the Blue module, which shows the strongest association with the biofilm phenotype, we found that this module is mainly enriched in stress response, chaperone, and other functional pathways. Additionally, the literature suggests that multiple hub genes are involved in the biofilm formation, virulence regulation, and bacteria pathogenicity (36–37).

### 3.5 PPI interaction networks between hub genes suggest novel interactions

Gene co-expression usually implies interactions or regulatory relationships among shared genes. Thus, we used the gene co-expression data obtained from WGCNA analysis to construct interaction networks. In order to compare with existing interaction data in public databases, we entered these genes to the STRING database, a current collection of known and predicted direct physical binding and indirect functionally related interactions between proteins/genes, and obtained their PPI (Protein-Protein Interaction) networks (38). Firstly, we performed PPI network mapping of hub genes within the three modules by gene co-expression relationships obtained from WGCNA, setting a weight threshold of the top 10%. The PPI networks constructed for the yellow and red modules’ hub genes demonstrated tight interactions, where functional enrichment was consistent with previous GO/KEGG analyses (Fig. S1). Interestingly, interaction of hub genes from the blue module sourced from the STRING database were significantly fewer than those from the WGCNA. Moreover, we found that the yellow module had a higher proportion of shared PPIs between both sources, accounting for 34% from WGCNA and 41.9% from STRING sources (Fig. 7D). In the red module, shared PPIs from WGCNA and STRING sources accounted for 54% and 32%, respectively (Fig. 7F). The blue module had the least shared PPIs, accounting for 5.2% from WGCNA and 56% from STRING sources (Fig. 7B). The variability of co-expression relationships in the blue module, as possibly indicated by STRING data, suggests there might be many new protein interactions within this module. Considering the blue module’s strongest association with the biofilm phenotype, members of the biofilm regulatory network might be located within this module, which are genes yet to be explored. Thus, through literature and public databases, we identified 6 genes encoding biofilm-associated proteins (42), also validated by biofilm-associated protein predictor. Based on the gene co-expression data from WGCNA, we obtained the PPI networks for these six biofilm-associated genes, respectively. By comparing gene members of each network and conducting overlapping analysis, we discovered 3 genes shared by all 6 networks, 7 shared by 5 networks, and 4 shared by 4 networks. Extracting the connections of these 14 shared genes with the 6 biofilm-associated genes to construct a PPI network, this network might represent a subnetwork of the biofilm formation pathway. These genes and their interactions await further experimental validation.

| Modules | Genes | Unannotated genes | Non protein-coding genes |
| --- | --- | --- | --- |
| Turquoise | 1,532 | 766 (50.0%) | 3 (0.2%) |
| Blue | 274 | 193 (70.4%) | 3 (1.1%) |
| Brown | 124 | 105 (84.7%) | 0 (0%) |
| Yellow | 76 | 41 (53.9%) | 0 (0%) |
| Grey | 67 | 42 (62.7%) | 0 (0%) |
| Green | 37 | 8 (21.6%) | 20 (54.1%) |
| Red | 34 | 15 (44.1%) | 0 (0%) |

### 3.6 The development of a comprehensive database of *Staphylococcus aureus*

The unassailable significance of making research data readily accessible cannot be overstated. To facilitate expeditious access to the data encompassed within this study for scholars in the field, we have established a designated database, named SAdb. Concurrently, we have amassed gene annotation information for *Staphylococcus aureus*, as well as transcriptomic data including both RNA-seq and microarray, alongside proteomic data. Utilizing this comprehensive suite of data, we performed differential expression analysis to obtain differentially expressed proteins and genes, conducted gene correlation analysis to obtain the correlations between genes, which were subsequently depicted via network visualization. These data and subsequent analyses constitute the four pivotal functional modules of the SAdb database. The “Gene” module is used to display gene annotation information on *Staphylococcus aureus*. The “Experiments” module collects experimental information on transcriptomics and proteomics. Clicking on a particular "Nr. total DEGs" entry will locate detailed information on DEGs under the corresponding experiment. In “Network” module, users can obtain the co-expression network of genes of interest through the search functionality. The “Results” module is used to display the data generated through this study. We hope this database could be an infrastructure for the biofilm research community.

## DISCUSSION

The exploration of biofilm formation by *Staphylococcus aureus* represents a pivotal area of inquiry within the domains of medical and biological research. Elucidating the underpinnings of biofilm development is instrumental in forging innovative therapeutic modalities aimed at dismantling the bulwarks of bacterial defiance, thereby furnishing novel methodologies for the amelioration of associated infections. In this study, we integrated transcriptomics projects from four experiments on *Staphylococcus aureus* biofilm phenotype, totaling 175 samples, with 85 samples in the bacterial motile state and 90 samples in the bacterial biofilm state. We hope to identify the bacterial molecular pathways affected in bacterial transition from motility to biofilm state as well as the regulatory networks of biofilm formation through comprehensive analysis.

Firstly, we conducted differential expression analysis and obtained 409 DEGs. Interestingly, GO and KEGG enrichment results revealed that only 32% (131/409) of the DEGs participated in enrichment, while the remaining 68% did not enrich any molecular pathways. This indicates that the annotation information of *Staphylococcus aureus* with public databases, including genes and functional pathways, is not comprehensive, greatly increasing analytical challenge. We used a differential fold ranking method to obtain the top 5 up-regulated and top 5 down-regulated genes. GO and KEGG analysis results showed that the urea metabolism pathway was affected, consistent with the enrichment results of feature genes selected using machine learning. The bacterial urea metabolism pathway, significant for environmental adaptability, virulence, and nitrogen utilization, intimates that during the transition from motile to biofilm state, pathogenicity are indeed affected, corroborating the postulation that “the infective prowess in the biofilm state is exponentially magnified in comparison to the motile state” (39).

The utilization of a differential fold ranking method, while rudimentary, overlooks certain genes with pivotal roles yet minute variations in expression, thus diverting from the research focus (40). GSEA emerges as a methodology concentrating on whether a preordained gene assembly exhibits systemic differential expression, proffering a panoramic view of biological processes, pathways, or functionalities (41). Despite the inconsequential expression alteration of solitary genes, the collective gene set may still command significant biological relevance. Nevertheless, the limited functional annotation information for *Staphylococcus aureus* curtails the availability of gene set data requisite for GSEA analysis. The application of machine learning analysis, The consideration of the variations in gene expression under different experimental conditions and the consistency of under the same experimental condition, to gene expression data often identifies important genes related to phenotype. Therefore, we used machine learning analysis to screen for feature genes, obtaining a total of 12. Contrasted with Top 12 genes, these feature genes enriched in more functional pathways, suggesting that machine learning analysis could capture a more comprehensive picture of pathway alterations. Additionally, several genes among the feature genes have been predicted to encode biofilm-associated proteins, consistent with literature reports (42), whereas fewer genes encoding biofilm-associated proteins were predicted among the Top genes. Comparing these two methods, we found that with sufficient samples of gene expression data, machine learning analysis may be more conducive to obtaining biologically meaningful genes than top DEGs.

Gene network analysis, as opposed to single gene analysis, can more comprehensively reveal the complexity and internal interactions of the system. Therefore, this study introduced WGCNA, hoping to obtain biologically meaningful gene modules. After obtaining gene modules, we performed hub gene analysis on three of the modules based on their relevance to experimental conditions and module membership composition. Functionally, these hub genes often enriched pathways associated with bacterial biofilm formation, virulence regulation, and pathogenicity, demonstrating the reliability of weighted gene co-expression network analysis in this study. In terms of the number of gene pairs, WGCNA obtained far more connections between genes than currently recorded in public databases. On one hand, this depicts that construction of a protein-protein interaction network based on hub genes has discovered novel interactions, a desired outcome. Yet, given that weighted gene co-expression network analysis fundamentally hinges on Pearson correlation to construct gene associations, it cannot circumvent the influence of indirect gene relationships, inevitably leading to numerous gene pairs with elevated correlation yet devoid of biological import. Considering direct gene relationships to construct a co-expression network epitomizes an efficacious stratagem for harvesting high-caliber gene pairs.

To facilitate the reuse and analysis of the data in this study by researchers in the field, and concurrently to archive data pertinent to *Staphylococcus aureus*, we have constructed a comprehensive database on *Staphylococcus aureus*. This database includes data generated from this study, as well as published transcriptomics and proteomics data. We hope this database can help researchers in the field better understand and analyze the genetic characteristics and pathogenesis of *Staphylococcus aureus*.

## 3. Materials and methods

### 3.1 Data collection, pre-processing and analysis

We downloaded raw sequencing data from ENA and SRA; evaluated the overall quality of raw sequencing reads using FastQC, then removed sequencing adapters and low-quality bases with Trim_galore, and performed transcript quantification with Salmon. After obtaining processed gene expression data, we integrated these data using the Combat function of the Sva R package and performed differential expression analysis, defining differentially expressed genes between experiments with cutoff values |log2 FC| > 1.0 (FC, fold change) and p-value <0.05. Additionally, we assessed the utility of common housekeeping genes and analyzed the function of Top DEGs.

### 3.2 Machine Learning Analysis

In the study, five popular ML algorithms including LR, XGBoost, SVM, KNN, and RF were used to analyze gene expression data to identify feature genes. All ML algorithms were implemented in Python and utilized grid search and 10-fold cross-validation for hyperparameter optimization.

### 3.3 WGCNA

Gene expression matrices were inputted to perform weighted gene co-expression network analysis using WGCNA (https://github.com/ShawnWx2019/WGCNA-shinyApp). This process included calculating the Pearson correlation matrix, adjacency matrix with power β = 3, and the final topological overlap matrix (TOM) based on normalized gene expression counts. We further filtered this TOM to exclude any samples containing weighted co-expressed <0.1 in all analyses.

### 3.4 Online Predictor

In this study, we used the BBSdb predictor, which we developed in previous research (), to predict obtained top DEGs, feature genes from machine learning model, and hub genes from important modules obtained through WGCNA, to discover potential biofilm-associated proteins.

### 3.5 2.5 Database Design and Implementation

SAdb is a relational database where all data are loaded into a MySQL database. The website’s frontend is coded using JavaScript and HTML, while the backend is coded in PHP to support queries to the MySQL database and provide a Representational State Transfer (REST) Application Programming Interface (API) for programmable access to our data. The AngularJS framework is used to connect the frontend and backend. Echarts.js and plotly.js are used for frontend visualizations.

## Supporting information

gene_expression

## Acknowledgements

The work was supported by grants from the NSFC (81871636 to EL).

